# Analytical methodology for unveiling the physical effect of detergents on the function of full-length membrane proteins using a single-molecule tracking system

**DOI:** 10.64898/2026.07.27.740883

**Authors:** Haruka Hoshi, Kenichi Kawano, Takahiro Fujiwara, Yousuke Yamaoka, Yusuke Kuroda, Kazuma Kurokawa, Aoi Taniguchi, Kenichi Nagase, Kiyosei Takasu, Katsumi Matsuzaki

## Abstract

G protein-coupled receptors (GPCRs) are important targets for drug discovery because they are the largest and most diverse family of membrane proteins in the human body. As exemplified by β_2_-adrenergic receptor (β_2_AR), which is a typical class-A GPCRs, they are prone to denaturation and inactivation after solubilization. Although diverse techniques have been developed, current structural and functional analyses are still limited to proteins with relatively stable and high expression. However, some mutant variants and misfolded proteins involved in diseases have extremely low expression levels and may be difficult to analyze. To overcome these limitations, we established a novel analytical platform for evaluating the ligand-binding ability of full-length β_2_AR as a model at the single-molecule level without purification and addressing challenges such as membrane proteins with low expression levels and structural instability. This method enables the direct use of unpurified receptors immediately after solubilization and allows us to use only a small amount of sample (∼10 ng) for observation and to distinguish between specific and nonspecific ligand binding by fitting. Furthermore, we unveiled the physical properties of detergents on the structural stability of solubilized receptors and found that the lateral pressure within detergent micelles affects ligand-binding ability. Detergents that provided a fluid microenvironment were able to maintain ligand-binding ability for several days even after solubilization; conversely, detergents that provided a rigid microenvironment caused the protein to lose its activity earlier. Our method could be a promising tool for the structural and functional analysis of membrane proteins untargeted until now.

**Graphical abstract:** 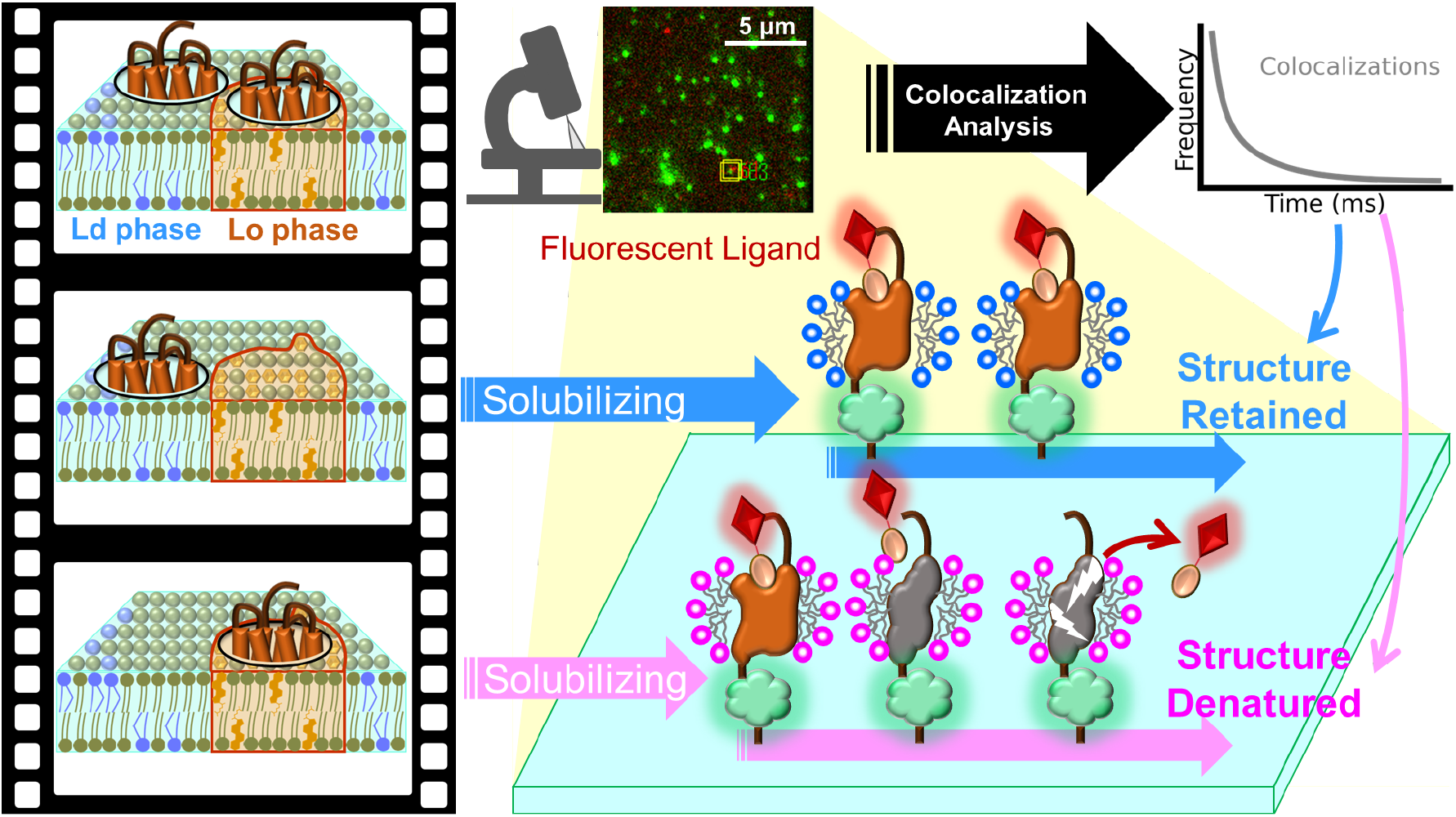

## Introduction

G protein-coupled receptors (GPCRs) are seven-transmembrane proteins that exhibit diverse physiological activities by binding to specific extracellular ligands, making them important targets for drug discovery^1^. As exemplified by β2-adrenergic receptor (β_2_AR), it is a typical class-A GPCR and has served as a model system since the early days of receptor research, whereas it is highly susceptible to denaturation and inactivation during and after solubilization^2^. Since the full-length cloning of β_2_AR in 1986 and the elucidation of its crystal structure in 2007, there has been rapid progress in the structural analysis of various GPCRs^3,4^. Cryo-electron microscopy is currently one of the most promising analytical techniques for the three-dimensional structure analysis of proteins^5^. Its practical application and widespread adoption have progressed rapidly, and it has become an indispensable technology for understanding molecules, including membrane proteins. In structural and functional studies using X-ray crystallography and cryo-electron microscopy, it is necessary to prepare large amounts of target proteins and isolate the receptors from biological membranes, removing unrelated contaminants as much as possible. The sample amount required for cryo-electron microscopy analysis is generally 3–30 µg, which is one-tenth the amount required for X-ray crystallography^6,7^. However, despite constant technological advancements make it possible that reduce the amount of protein needed for analysis, there is still a tendency to target structurally stable proteins with relatively high expression levels for analysis. This means that some mutant variants and misfolded membrane proteins involved in diseases have extremely low expression levels and may be impossible to analyze. Therefore, developing analytical platforms for single-molecule functional analysis or novel ligand screening against rare membrane proteins without purification is an urgent task. As the first step to overcome this demand, in this study, we developed a single-molecule observation system and analytical methodology for evaluating the ligand-binding ability of full-length β_2_AR as a model over several days at 25°C without any modification to the core sequence by employing total internal reflection fluorescence microscopy (TIRFM). This method allows us to use unpurified β_2_AR immediately after solubilization to measure the interaction with its ligand using a small sample volume (∼30 µL containing ∼10 ng of proteins of interest) and to distinguish between nonspecific and specific ligand binding by fitting. Because TIRFM is a technique that utilizes evanescent waves penetrating only up to 100 nm from the glass substrate, enabling the highly sensitive detection of faint fluorescence signals from a small amount of samples on the immediate vicinity of the glass surface, it is suitable for a single-molecule tracking system of the colocalization between solubilized β_2_AR and its fluorescent ligand.

In general, solubilized GPCRs tend to lose their native structures and functions inherent to biological membranes, and retaining their stability remains a major challenge. Approaches that modify the membrane proteins themselves to improve their stability, such as mutation introduction into the core sequence^8^, chimera formation with fusion partner proteins in the 3^rd^ intracellular loop^2,4^, and C-terminal truncation^9^, have led to the development of a variety of agents with the aim of solubilizing the target to preserve its native state for as long as possible. In addition to commonly used detergents, numerous alternative methods have been reported, including polymer nanodiscs, membrane scaffold protein nanodiscs, and artificial liposomes^10^. Although diverse approaches for membrane protein solubilization exist, there is still room for investigation regarding the specific microenvironments provided by different detergent micelles to GPCR targets and how differences in the physical properties affect the ligand-binding ability and stability of solubilized GPCRs. In this study, three types of detergents were selected to solubilize the target β_2_AR: (*i*) mixed micelles of 1-oleoyl-2-cholyl-*sn*-glycelo-3-phosphocholine and 1-oleoyl-2-(3, 12-disulfo)deoxycholyl-*sn*-glycelo-3-phosphocholine (hereafter, cholyl-PC + di-sulf(D)), (*ii*) lauryl maltose neopentyl glycol (LMNG), and (*iii*) *n*-dodecyl-β-D-maltoside (DDM)^11–13^. These detergents offer advantages such as the convenience of directly solubilizing proteins from biomembranes for immediate analysis and the relatively low cost of large-scale synthesis. LMNG and DDM are widely used in structural analyses of membrane proteins^14–16^. In a previous study, these detergents demonstrated a good balance of high solubilization efficiency and low denaturing effects on bacteriorhodopsin, a proton channel, compared to other detergents^11^. Structurally, cholyl-PC + di-sulf(D) possesses a phospholipid head group, an oleoyl group, and a cholic acid-derived steroid backbone, whereas LMNG and DDM are composed of alkyl glycosides (Fig. 1). Because cholyl-PC + di-sulf(D) mimics the lipid components of biological membranes, they have a compact hydrophilic group and features a structurally rigid steroid backbone; therefore, it is expected to exhibit different physical properties from LMNG and DDM. The concept of this study was to establish an evaluation system to achieve a simple comparison of the effects of detergents on the ligand-binding stability of solubilized GPCRs under unpurified conditions. Based on these results, we discuss the differences in receptor stability induced by the physical properties of detergent micelles.

**Fig. 1.**
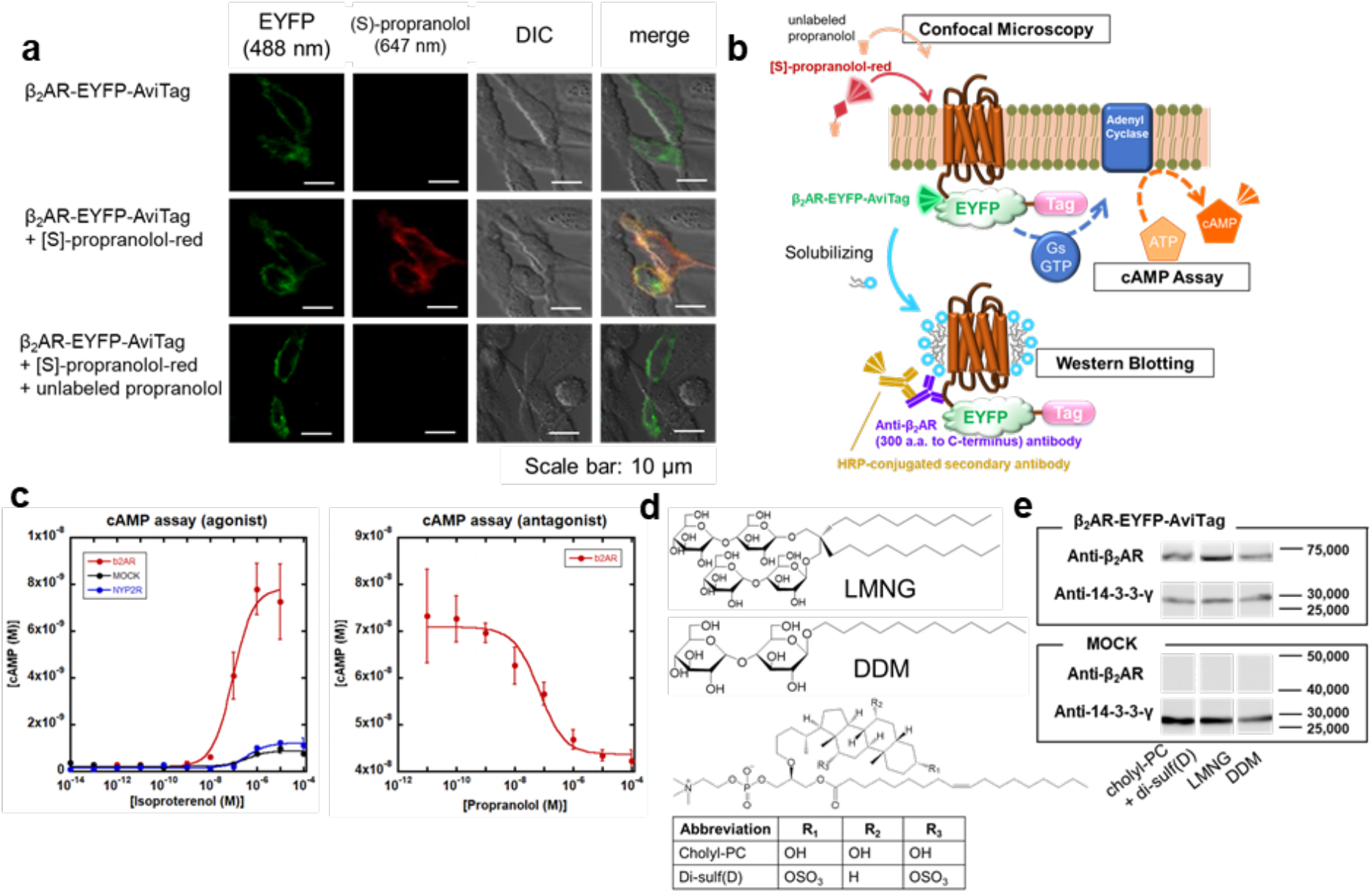
β_2_AR-EYFP-AviTag expression and cAMP responses in HEK293 cells, and immunoblotting analysis after solubilization. **(a)** Confocal microscopy images of β_2_AR-EYFP-AviTag. The top and middle images were acquired in the same fields of HEK293T cells before and 20 min after the addition of 50 nM (S)-propranolol-red. The bottom images were obtained 20 min after the addition of 50 nM (S)-propranolol-red and 10 µM unlabeled propranolol. **(b)** Schematic representation of cAMP signaling and western blotting using anti-β_2_AR antibody recognizing the C-terminal region from 300 a.a.. (c) cAMP response curves in HEK293T cells upon treatment with isoproterenol or propranolol (mean ± S.D., *n* = 3). **(d)** Structural formulas of detergents: cholyl-PC, di-sulf(D), LMNG, and DDM. (e) Western blotting of β_2_AR and 14-3-3-γ.

## Materials and methods

### Preparation of β_2_AR and NPY_2_R constructs

The amino acid sequences of all constructs are shown in Fig. S1. All genes were cloned into the pcDNA 3.1 vector (Thermo Fisher Scientific, MA, Waltham). Plasmid DNA encoding E3-β_2_AR-EYFP^17^ was used for gene modification. The E3 tag was removed using the HindIII restriction enzyme, and immobilization tags were introduced at the C-terminal site downstream of EYFP using the BsrGI and XhoI restriction enzymes. For NPY_2_R, an immobilization tag was introduced at the C-terminus of the hNPY_2_R-EYFP construct^8^. The SBP and AviTag sequences were designed with reference to the RCSB (PDB: 4JO6) and previous studies^18,19^, respectively.

### Confocal microscopy observation

HEK293T cells (cell number RCB2202), which was provided by the RIKEN BRC CELL BANK (Tsukuba, Japan), were cultured in D-MEM (Nacalai Tesque, Kyoto, Japan) supplemented with 10% heat-inactivated fetal bovine serum at 37°C under 5% CO_2_. HEK293T cells were seeded in a 35 mm glass-bottom dish (Iwaki, Tokyo, Japan) at a density of 3 × 10^5^ cells/dish. After 24 h of incubation, the medium was replaced with serum-free medium. A mixture containing 4 µL of Lipofectamine LTX Reagent (Thermo Fisher Scientific), 2 µg of pDNA, and 400 µL of Opti-MEM was then added to the dish, and the cells were incubated for 6 h. After post-transfection, the medium was replaced with serum-containing medium, and the cells were used for experiments 18–42 h later. The expression of β_2_AR-EYFP or NPY_2_R-EYFP constructs transiently expressed in HEK293T cells was observed in phosphate-buffered saline PBS(+) [137 mM NaCl, 8.1 mM Na_2_HPO_4_, 2.68 mM KCl, 1.47 mM KH_2_PO_4_, 0.33 mM MgCl_2_, and 0.90 mM CaCl_2_, pH 7.4] using a confocal laser scanning microscope, A1R-MP (Nikon, Tokyo, Japan). Ligand binding was observed after incubating cells with 50 nM (S)-propranolol-red (CA200689) (Hello Bio, Bristol, UK), 100 nM TMR-NPY^11^, or 1 µM unlabeled propranolol at 25°C.

### cAMP assay

The cAMP response of HEK293T cells transfected with β_2_AR-EYFP-AviTag or empty vector (MOCK) was measured using the AlphaScreen cAMP assay kit (PerkinElmer, Shelton, CT) according to the manufacturer’s instructions with partial modifications. The cells transfected using a reagent, linear polyethylenimine (Tokyo Chemical Industry, Tokyo, Japan), were seeded into a 384-well plate at a density of 2,500 cells/well and stimulated with ligands for 30 min at 37°C. The donor beads (33.4 µg/mL) with biotinylated cAMP (16.7 nM) and acceptor beads (200 µg/mL) were sequentially added to the wells and incubated for 1 h at 25 °C under light-blocking conditions. The signal was measured using a plate reader, EnSight (Levvity, Waltham, MA), and then converted to cAMP concentrations based on a standard curve. The EC_50_ (or IC50) value was obtained by fitting sigmoidal curves using the following equation, as previously described^20^:

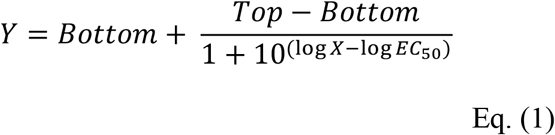

where *Y* and *X* denote the concentration of cAMP and ligands, *Top* and *Bottom* indicate the top and bottom of the curve asymptote, respectively.

### Preparation of detergents

Cholyl-PC and di-sulf(D) were synthesized as described in a previous study^11^. LMNG and DDM were purchased from Anatrace (Maumee, OH) and Nacalai Tesque, respectively.

### Solubilization of GPCRs

HEK293T cells were seeded in a 100 mm dish at a density of 2.4 × 10^6^ cells/dish. Transfection was performed using a mixture of 32 µL LTX, 16 µg DNA, and 3.2 mL Opti-MEM. After 18–42 h, the cells were washed with 10 mL PBS(–). Subsequently, 1 mL/dish of detergents in PBS(–) at final concentrations (1 mM cholyl-PC, 0.5% LMNG, or 1% DDM) containing 1 tablet/10 mL protease inhibitor cocktail, cOmplete^TM^ (Roche, Mannheim, Germany), was incubated with cells at 25°C for 90 min. The cells were gently detached using a scraper and incubated for an additional 30 min at 25°C. The supernatant was collected after centrifugation at 13,800 × *g* for 5 min at 25°C. Di-sulf(D) at a final concentration of 1 mM was mixed with the cholyl-PC solution and incubated for an additional 1 h at 25°C.

### Western blotting

Western blotting was performed as described study^21,22^. Briefly, solubilized proteins were mixed with SDS sample buffer and separated by SDS-polyacrylamide gel electrophoresis (10%) at 30 mA for 75 min. Proteins on the gel were transferred onto a membrane (WSE-4051) (Atto, Tokyo, Japan) at 2.5 V for 30 min. After blocked with 5% (w/v) skim milk for 1 hr, the membrane was incubated with the primary antibodies, rabbit anti-human β_2_AR antibody (bsm-52880r) (Bioss Antibodies, Woburn, MA) or anti-14-3-3 gamma antibody (GTX134246) (GeneTex, Irvine, CA) at 1:1,000 and 1:2,000 dilutions, respectively. The secondary antibody, HRP-conjugated goat anti-rabbit IgG(H+L) (SA00001-2) (Proteintech, Rosemont, IL) was used at a 1:2,000 dilution. Protein bands were then detected by using a chemiluminescent substrate SuperSignal West Pico PLUS (Thermo Fisher Scientific) and an imaging system, ImageQuant LAS 4010 (Cytiva, Marlborough, MA).

### Glass substrate preparation and TIRFM observation

Before TIRFM observation, ultrafiltration of solubilized protein was performed using an ultra-centrifugal filter, 30k MWCO Amicon UFC5030 (Millipore, Burlington, MA) at 14,000 × *g* and 25°C for 5 min to improve biotin-labeling efficiency at the following step. AviTag biotinylation was performed using a BirA biotin-protein ligase standard reaction kit (Avidity, Aurora, CO) by adding 2.5 µL of BirA to 250 µL of the solubilized solution, followed by overnight incubation at 30°C. To remove unreacted biotin remaining in the solution, centrifugation for 5 mins using the 30k MWCO filter was repeated five times. The sample was then added to the observation chamber, which was composed of a slide glass (25.7 × 75.7 mm (1.0–1.2-mm thickness) (Matsunami Glass, Osaka, Japan) with two 2-mm diameter holes drilled in the center, a silicone spacer (0.5-mm thickness) (As One, Osaka, Japan), and a cover glass (24 × 60 mm, Matsunami Glass) as following the previous studies^11,21,23^. The chamber was washed with each detergent solution and then filled with a solution of 50-fold diluted solubilized GPCRs (30 µL)^23,24^. After incubation for 10 min, unreacted GPCRs were removed by washing, and GPCR immobilization was confirmed by observing EYFP fluorescent spots using TIRFM. Subsequently, 1 nM of a fluorescent ligand was injected into the chamber and incubated for 2 h for sufficient interaction with GPCRs. A low concentration of the fluorescent ligand (1 nM) was necessary to achieve a high S/N ratio in the single-molecule measurement. The fluorescent spots of EYFP and ligands were recorded at two independent channels at the same time for 1,000 frames at 60 fps as following the previous _studies11,25,26._

### Fluorescent spot tracking and colocalization analysis

Fluorescent spot tracking and colocalization analyses were conducted based on the methods developed previously^26,27^. To account for the transient blinking of fluorescent molecules, if a maximum of five dark frames was observed in the tracking process, these frames were treated as a single continuous trajectory traced by the diffusion of the same bright spot; however, if the bright spot disappeared for six frames or more, the trajectory was terminated. In the colocalization analysis, the distance threshold between the fluorescent spots of solubilized GPCRs and ligands was set to 450 nm, as shown in Figure 3a. Freely diffusing ligands passed through the evanescent field without binding to the receptors; therefore, to eliminate the influence of these transient and noise-derived signals, colocalization events with a dwell time of 7 frames or shorter (< 120 ms) were excluded from the subsequent curve fitting analysis. The frequency of colocalization was plotted as a function of the duration time and fitted with single-or double-exponential decay equations, as shown below^25^ under the assumption of the previous model^28^:

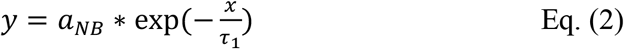

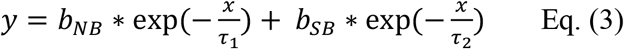

where *a*_NB_ denotes the amplitude for nonspecific binding (NB) of (S)-propranolol-red against NPY_2_R-EYFP-AviTag, and *b*_NS_ and *b*_SB_ indicate the amplitudes for NB and specific binding (SB) of (S)-propranolol-red against β_2_AR-EYFP-AviTag, respectively, *τ*_1_ and *τ*_2_ are the time constants for NB and SB, respectively. Here, it was assumed that the *τ*_2_ value for SB is larger than the *τ*_1_ value for NB because *τ* is defined as being inversely proportional to the reciprocal of the dissociation rate constant, *k*_off_ (*τ* = 1/*k*_off_), indicating that a larger *τ* value means a longer binding time for the ligand to dissociate due to SB. To determine the *b*_SB_ value, we first estimated the *τ*_1_ value for each detergent condition by fitting the colocalization frequency curve for NPY_2_R-EYFP-AviTag + (S)-propranolol-red with Eq. (2). Next, we fixed the *τ*_2_ value at 300, because when the decay curves for β_2_AR-EYFP-AviTag + (S)-propranolol-red were fitted with Eq. (3) using *τ*_1_ as a constant and *τ*_2_ as a variable, the *τ*_2_ value consistently converged to approximately 300 ms regardless of the type of detergents. Finally, using *τ*_1_ determined from Eq. (2) and *τ*_2_ fixed at 300, we fitted the colocalization frequency curve for β_2_AR-EYFP-AviTag + (S)-propranolol-red with Eq. (3) to determine the *b*_SB_ value.

### Calculation of GP values

Detergents at the same concentrations described above were incubated with 1 µM Laurdan for 2 h. The fluorescence spectra of Laurdan were measured at 0, 24, and 48 h after preparation at 25°C using a spectrofluorometer, RF-6000 (Shimadzu, Kyoto, Japan) with excitation at 350 nm and emission at 400–600 nm. After subtracting the background signal, the GP value was calculated based on the fluorescence intensities at 440 nm (*I*_440_) and 490 nm (*I*_490_) according to the following equation:

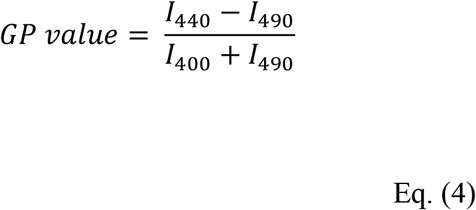

CHS was mixed with LMNG or DDM at a final concentration of 0.05%. Liposomes were prepared with a lipid composition of 25% 1-palmitoyl-2-oleoyl-*sn*-glycero-3-phosphatidylcholine (POPC), 22% 1-palmitoyl-2-oleoyl-*sn*-glycero-3-phosphatidylethanolamine (POPE), 10% 1-palmitoyl-2-oleoyl-*sn*-glycero-3-phosphatidylserine (POPS), 18% sphingomyelin (SM), and 25% cholesterol (mol%) in Bis-Tris buffer, as described in a previous study^29^. Laurdan was incubated with liposomes for 2 h at 25°C for labeling the vesicles. The particle size was measured using dynamic light scattering (DLS) with a Zetasizer Nano ZS (Malvern Panalytical, Malvern, UK). HEK293T cells were seeded in a 60 mm dish at a density of 3.0 × 10^5^ cells/dish. After 2 days of culture, 1 µM Laurdan in PBS(+) was added to the cells and incubated for 2 h at 37°C in a 5% CO_2_ incubator. Subsequently, the stained cells were detached using trypsin–EDTA and used for measuring the fluorescence spectrum at a concentration of 1.4 × 10^6^ cells/mL at 25°C.

Refer to the *Supplemental Materials and Methods* section for FRET and FCS measurements.

## Results

### β_2_AR construct design and its functional activity in living cells

β_2_AR was selected as a model GPCR for analyzing the microenvironmental effects of detergents on its functional activity. The C-terminus of human β_2_AR was fused with an enhanced yellow fluorescent protein (EYFP) and an immobilization tag for fluorescent microscopic observation (Fig. S1). The C-terminal fusion construct was designed to ensure that the target membrane protein was present in its full-length form. AviTag^TM^ (GLNDIFEAQKIEWHE) and streptavidin-binding peptide (SBP) (MDEKTTGWRGGHVVEGLAGELEQLRARLEHHPQGQREP) were chosen as the immobilization tags on the glass surface because of their strong interactions with avidin^30^. The former has a lysine residue within the sequence for biotinylation by a biotin ligase, BirA, and tightly interacts with avidin (dissociation constant: ∼10^-15^ M)^19^, and the latter is an affinity purification tag with an equilibrium dissociation constant of 2.5 nM for streptavidin^18^. Before the solubilization experiments, we confirmed the expression of the representative construct (β_2_AR-EYFP-AviTag) and its ligand-binding ability in human embryonic kidney 293T (HEK293T) cells. EYFP fluorescence for β_2_AR-EYFP-AviTag was observed on the cell membrane and colocalized with that of its commercially available fluorescent antagonist, (S)-propranolol-red, using confocal microscopy (Fig. 1a), suggesting that the β_2_AR construct has ligand-binding ability. In the presence of an excess amount of unlabeled propranolol, the fluorescence signal of (S)-propranolol-red disappeared from the membrane, indicating that the fluorescent antagonist specifically bound to the ligand pocket of β_2_AR^31^.

We also confirmed that β_2_AR-EYFP-AviTag has functional activity in cyclic adenosine 5’-monophosphate (cAMP) signaling against ligand stimulation in living cells (Fig. 1b). β_2_AR-EYFP-AviTag exhibited a dose-dependent cAMP response upon treatment with its agonist (isoproterenol) with an EC_50_ value of 9.3 × 10⁻⁸ M (Fig. 1c), which is similar to the reported EC_50_ value^32^. In contrast, a slight cAMP response was observed for endogenous β_2_AR (empty vector-transfected MOCK cells) at 3.7 × 10⁻⁷ M (Fig. 1c). Notably, β_2_AR-EYFP-AviTag exhibited approximately 7–8-fold higher [cAMP (M)] than endogenous β_2_AR, demonstrating that cAMP production is proportional to receptor expression levels. We also prepared a control class-A GPCR, the human neuropeptide Y2 receptor, with the same fusion constructs (NPY_2_R-EYFP-AviTag and NPY_2_R-EYFP-SBP) (Fig. S1)^11^. The expression of NPY_2_R-EYFP-AviTag and its binding to the original ligand peptide (NPY) were confirmed in HEK293T cells (Fig. S2), its cAMP response against isoproterenol stimulation was negligible, similar to that of MOCK (EC_50_:4.1 × 10⁻⁷ M) (Fig. 1c), indicating that NPY_2_R-EYFP-AviTag does not respond to β_2_AR ligand stimulation. In contrast, β_2_AR-EYFP-AviTag also responded to propranolol stimulation followed by forskolin treatment and reduced cAMP levels in a dose-dependent manner (Fig. 1c) (IC_50_ of 6.9 × 10⁻⁸ M), similar to the reported value^33^, suggesting that our designed β_2_AR construct possesses both ligand-binding and signal transducing abilities.

The chemical structures of the detergents, cholyl-PC + di-sulf(D), LMNG, and DDM^11–13^, used in this study are shown in Figure 1d. β_2_AR-EYFP-AviTag expressed in HEK293T cells was solubilized using these detergents and detected near the expected molecular weight of approximately 75,000 by western blotting using an anti-β_2_AR C-terminus antibody (Fig. 1b and e). The housekeeping protein 14-3-3-γ was detected at similar expression levels among these detergents near the expected molecular weight of approximately 28,000. In contrast, the immunoblotting band of endogenous β_2_AR in MOCK cells was below the detection limit (Fig. 1e), indicating that the influence of endogenous β_2_AR can be disregarded in the following experiments.

### Optimization of β_2_AR immobilization conditions for TIRFM observation

To establish a single-molecule observation system enabling the analysis of the ligand-binding ability of solubilized β_2_AR immobilized onto glass substrates under unpurified conditions, we optimized the immobilization method to achieve the best signal-to-noise (S/N) ratio. While TIRFM is a powerful technique capable of capturing single-molecule fluorescent spots with high spatial resolution, improving the S/N ratio makes it possible to minimize false-positive signals. The biotin–avidin interaction is widely used for the immobilization and purification of proteins owing to its high affinity and ease of labeling^34^. In this study, we compared two types of immobilization tags (AviTag vs. SBP) and two types of coating materials (bovine serum albumin: BSA vs. polyethylene glycol: PEG) to identify the optimal combination (Fig. 2). In the first step, the glass substrate was coated with BSA-biotin directly or amino-functionalized with a silane coupling agent and then reacted with a mixture of methoxy PEG succinimidyl valerate (mPEG-SVA with an average MW of 5,000) and its biotin derivative (PEG-biotin) (Fig. 2a), following the reported methods^23,24^. While BSA allows for simple coating through physical adsorption onto glass, the method using PEG-biotin is thought to enable more robust immobilization through covalent chemical modification achieved by applying a silane coupling agent to the glass substrate. In the second step, β_2_AR-EYFP-AviTag or β_2_AR-EYFP-SBP was immobilized onto the glass substrate through a strong interaction with Neutravidin (streptavidin was also examined, but similar results were obtained; data not shown) (Fig. 2a). AviTag was biotinylated by BirA in advance^18,19^. Using these four combinations, we observed fluorescent signals derived from solubilized β_2_AR-EYFP in cholyl-PC + di-sulf (D) micelles using TIRFM. The captured images of β_2_AR-EYFP with line profiles, as shown in green, are presented in Figure 2b. The two conditions employing the SBP-tag showed several identifiable fluorescent spots, suggesting low immobilization efficiency under the present conditions. Stray light from adjacent spots or background signals from nonspecific adsorption on the substrate may interfere with the distinct identification of individual immobilized β_2_AR spots. In contrast, the other two conditions utilizing AviTag showed clear fluorescence signals; in particular, the combination with PEG-biotin coating ((2) and (3)) achieved a relatively higher S/N ratio by minimizing the background signals to near zero (Fig. 2b). Moreover, the best average number of spots/frame was detected as 65±3 under the same conditions, except for the (1) and (3) combinations, which had the lowest S/N ratio (Fig. 2b). This provides ideal conditions for single-molecule observation, allowing for the identification of individual spots. Based on these findings, we adopted a combination of PEG-biotin coating and AviTag ((2) and (3)) as a suitable system for single-molecule observation.

**Fig. 2.**
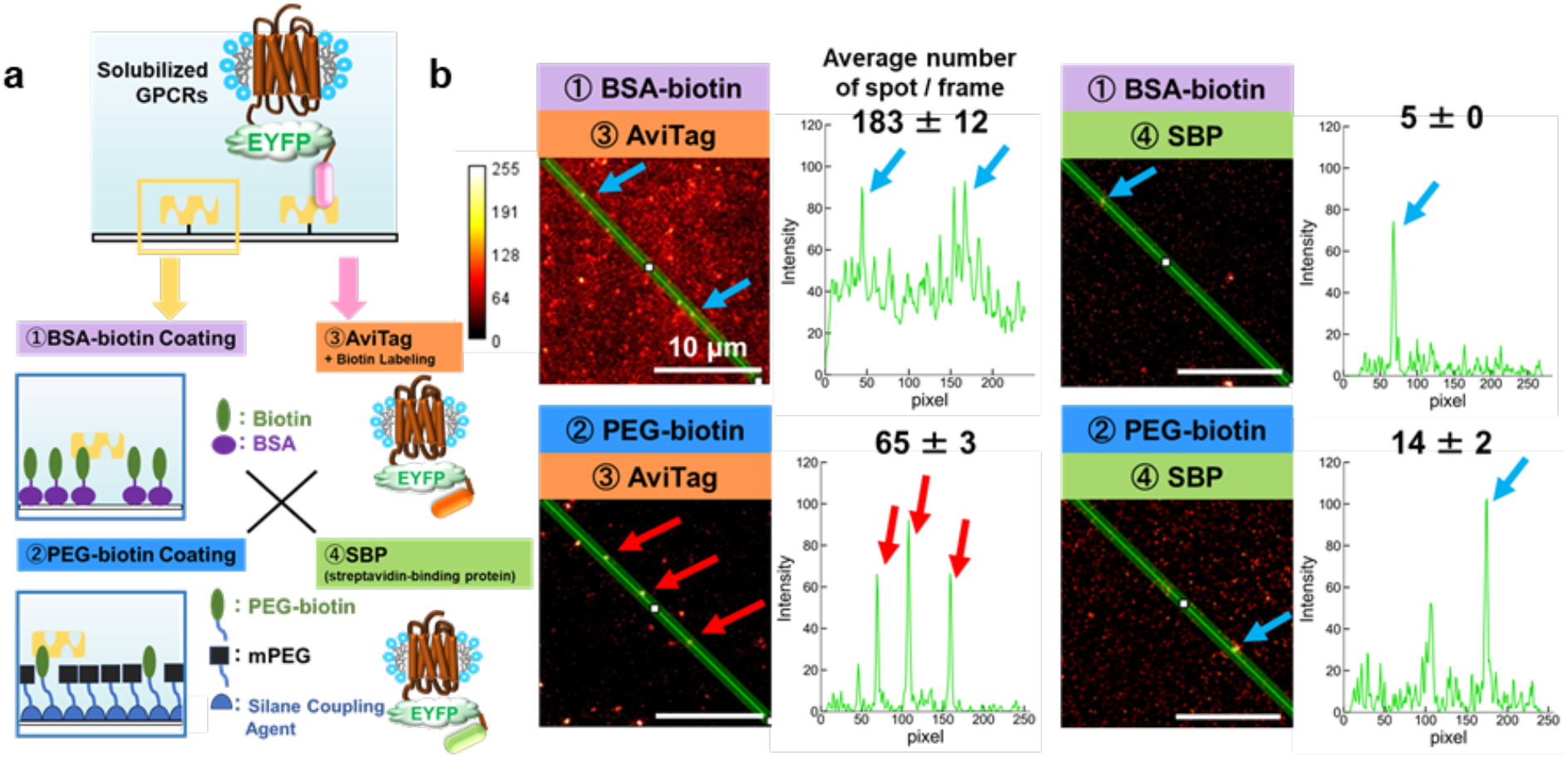
Optimized conditions for β_2_AR immobilization onto glass substrates for TIRFM observation. **(a)** Schematic of the immobilization conditions examined in this study. **(b)** TIRFM images (left) and fluorescence intensity line profiles (right) of β_2_AR immobilized under each condition. The images are 8-bit and pseudo-colored according to the color bar shown on the left. The fluorescence intensity profile along the green line in the image is plotted from the upper left to the lower right. The arrows in the figure indicate the intersection points of the green line and fluorescent spots. The average number of spots/frame is shown above each line profile.

**Fig. 3.**
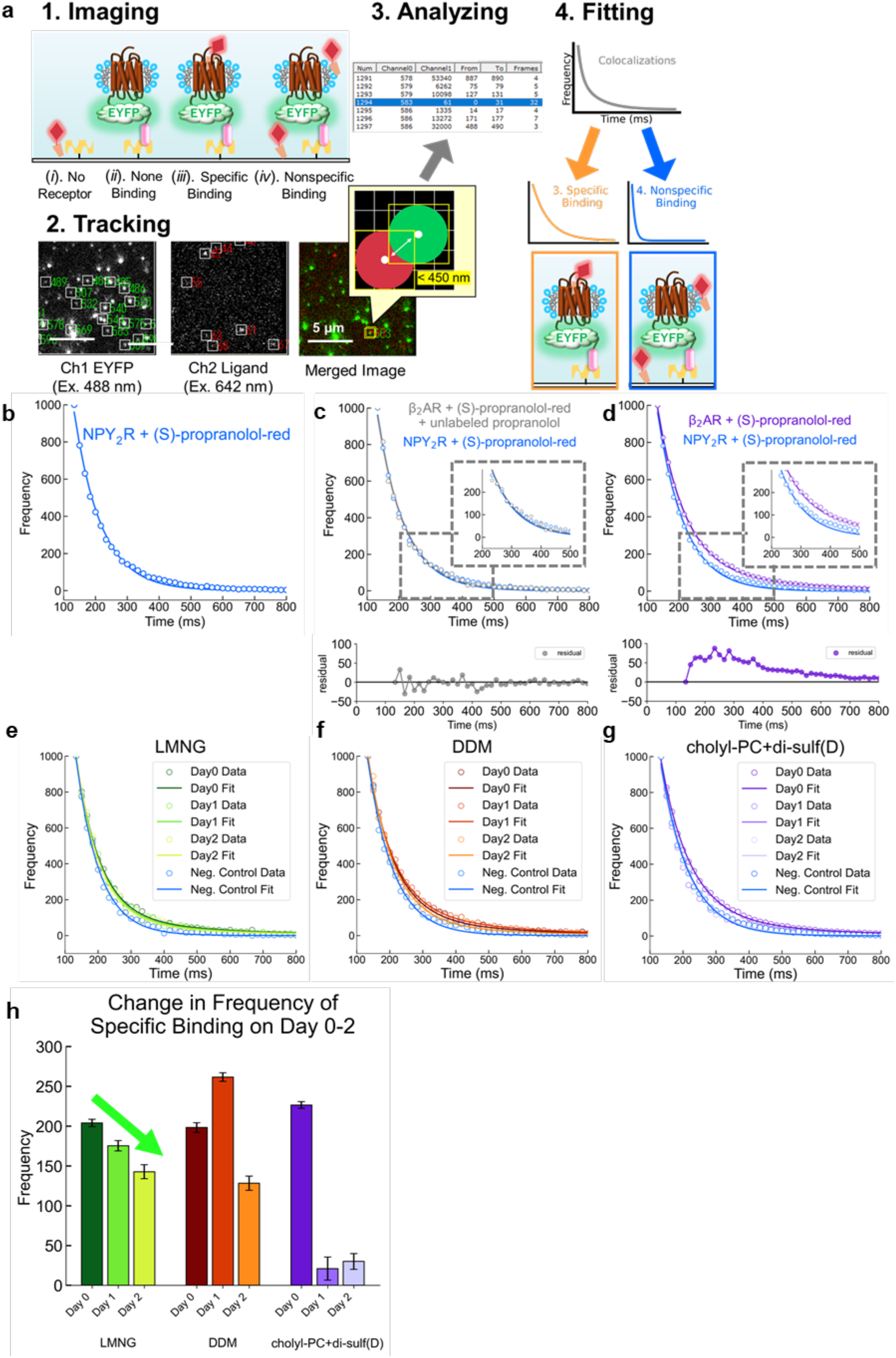
Co-localization analysis of solubilized β_2_AR and a fluorescent antagonist for specific ligand-binding analysis. **(a)** Overview of the co-localization analysis and ligand-binding evaluation. The colocalization in this study was defined as occurring when the distance between two-color fluorescent spots approached each other within 450 nm, about twice the radius of Airy Disk based on the Rayleigh criterion. A brief procedure for imaging and tracking of fluorescent spots for solubilized β_2_AR-EYFP-AviTag and (S)-propranolol-red captured by TIRFM, and fitting for analyzing the specific ligand-binding component are shown in this panel. **(b–d)** Colocalization frequency plotted as a function of the duration time (ms) for NPY_2_R + (S)-propranolol-red, β_2_AR + (S)-propranolol-red + excess unlabeled propranolol, and β_2_AR + (S)-propranolol-red. The residuals are shown in panels **(c)** and **(d)**. (S)-Propranolol-red and unlabeled propranolol were used at final concentrations of 1 nM and 1 µM, respectively. **(e–g)** Time-lapse change in the decay curves for β_2_AR + (S)-propranolol-red in each detergent on Days 0,1, and 2. **(h)** Transitions in the specific ligand-binding frequency for β_2_ARs solubilized in each detergent, which was calculated based on Eq. (3) (mean ± S.D., *n* = 3).

### Time-lapse ligand-binding analysis of solubilized β_2_AR by TIRFM

Based on the above-mentioned optimized conditions, we developed a method for analyzing the ligand-binding ability of target membrane proteins. This method relies on single-molecule tracking and colocalization analysis^26,27^ of solubilized β_2_AR-EYFP-AviTag in micelles and its fluorescent antagonist, (S)-propranolol-red, and allowed us to evaluate the detergents suitable for the target membrane proteins. Unlike existing methods, this approach does not require the purification of membrane proteins; therefore, its greatest advantage is that it allows for the analysis of their functional activities at a small volume (∼30 µL containing ∼10 ng of proteins) using fresh samples immediately after solubilization. We evaluated how long β_2_AR in micelles could retain its ligand-binding ability over several days at 25°C. Notably, EYFP (Ex. 488 nm) and (S)-propranolol-red (Ex. 642 nm) were selected as dyes for this dual-imaging application because they do not undergo fluorescence resonance energy transfer (FRET).

Solubilized β_2_AR-EYFP-AviTag was immobilized onto the PEG-biotin-coated glass substrate, and (S)-propranolol-red was added. Subsequently, the fluorescent spot images of EYFP and (S)-propranolol-red were simultaneously recorded in two independent channels for 1000 frames at 60 fps (Fig. 3a). Theoretically, the observed fluorescent spots of β_2_AR-EYFP-AviTag and (S)-propranolol-red can be classified into the following four categories: (*i*) ligands bound non-specifically to the glass substrate, (*ii*) receptors without bound ligands, (*iii*) receptors with ligands bound specifically to the binding site, and (*iv*) receptors with ligand bound non-specifically to micelles or the surrounding glass surface (Fig. 3a). Categories (*i*) and (*ii*), which show single-color spots, can be excluded from the colocalization analysis. Categories (*iii*) and (*iv*), which include dual-color spots, were deconvoluted into specific and non-specific binding components based on their duration lifetime (*τ*) in the colocalization analysis (Fig. 3a). In this study, “colocalization” was defined as occurring when the distance between two-color fluorescent spots approached each other within 450 nm (Fig. 3a), about twice the radius of Airy Disk based on the Rayleigh criterion, considering the EYFP fluorescence wavelength of 525 nm and the numerical aperture of 1.49 for the oil immersion objective lens^35^. The decay curve was fitted using a single-exponential Eq. (2) or the double-exponential Eq. (3) (refer to the *Materials and methods* section) to obtain nonspecific and specific binding components, respectively. The fitting results are summarized in Table 1, and the latter factor is shown in Figure 3h.

We will explain the details using actual experimental data. First, the nonspecific binding component of (S)-propranolol-red against micelles (category (*iv*)) (Fig. 3a) was estimated using the negative control NPY_2_R-EYFP-AviTag. This is because propranolol (a β_2_AR antagonist) does not interact with the peptide ligand-binding pocket of NPY_2_R. The colocalization frequency curve for solubilized NPY_2_R-EYFP-AviTag in a mixture of cholyl-PC + di-sulf(D) and (S)-propranolol-red (Fig. 3b) was fitted with Eq. (2), and the *τ*_1_ value of nonspecific binding was determined to be 85 ms (Table 1). Here, the y-axis of the decay curve was normalized to 1000 before fitting. Importantly, the decay curve of NPY_2_R-EYFP-AviTag perfectly overlapped with that of β_2_AR-EYFP-AviTag + (S)-propranolol-red in the presence of an excess amount of unlabeled propranolol (categories (*iv*) vs. (*iv*)) (Fig. 3c), demonstrating that the nonspecific binding component was accurately estimated. As proof of that, the residual between the two curves exhibited almost zero (Fig. 3c), In contrast, the decay curve of the nonspecific binding was not overlapped with that of β_2_AR-EYFP-AviTag + (S)-propranolol-red in the absence of unlabeled propranolol (categories (*iv*) vs. (*iii*)+(*iv*)) (Fig. 3d), and the residual between the two curves obviously remained (Fig. 3d). These data demonstrate that the nonspecific and specific binding components were properly estimated using this approach.

Using the obtained *τ*_1_ value as a constant, we obtained the *b*_SB_ value (the specific binding parameter) of 227 ± 4 for β_2_AR-EYFP-AviTag in a mixture of cholyl-PC + di-sulf(D) on Day 0 by fitting using Eq. (3) (Fig. 3e) (Table 1). The same procedure was adapted for solubilized β_2_AR-EYFP-AviTag in LMNG and DDM micelles (Fig. 3f and g). On Day 0, immediately after solubilization, the *b*_SB_ values were similar among all detergents, indicating that specific ligand binding occurred at comparable levels (Fig. 3h, Table 1). To examine whether solubilized β_2_AR retains its ligand-binding ability over days at 25 °C, we tracked the temporal changes in ligand binding up to two days post-solubilization. β_2_AR-EYFP-AviTag on Days 1 and 2 exhibited lower *b*_SB_ values, which were similar to that of 26 ± 11 for β_2_AR-EYFP-AviTag in the presence of an excess amount of unlabeled propranolol (Fig. 3c) (Table 1). Although the *b*_SB_ values for solubilized β_2_AR-EYFP-AviTag in LMNG or DDM micelles gradually decreased, they remained at relatively higher levels (Fig. 3h). These results indicate that the ligand-binding ability of β_2_AR is highly affected by detergents. To reveal the underlying reason for these phenomena, we evaluated the physical properties of micelles formed by individual detergents.

### Investigation of the physical properties of micelles

The physical properties of micelles are required to separate the surficial and internal factors of micelles; that is, the former assumes micelle or protein aggregations, causing spatial ligand access restrictions to the binding pocket, and the latter assumes a high lateral pressure within micelles, causing a loss of the three-dimensional structure for ligand binding. Based on these assumptions, we investigated and found that micelle or protein aggregations did not occur (Figs. S3 and S4), rather the lateral pressure within the micelles influences the activity of solubilized β_2_AR (Fig. 4).

**Fig. 4.**
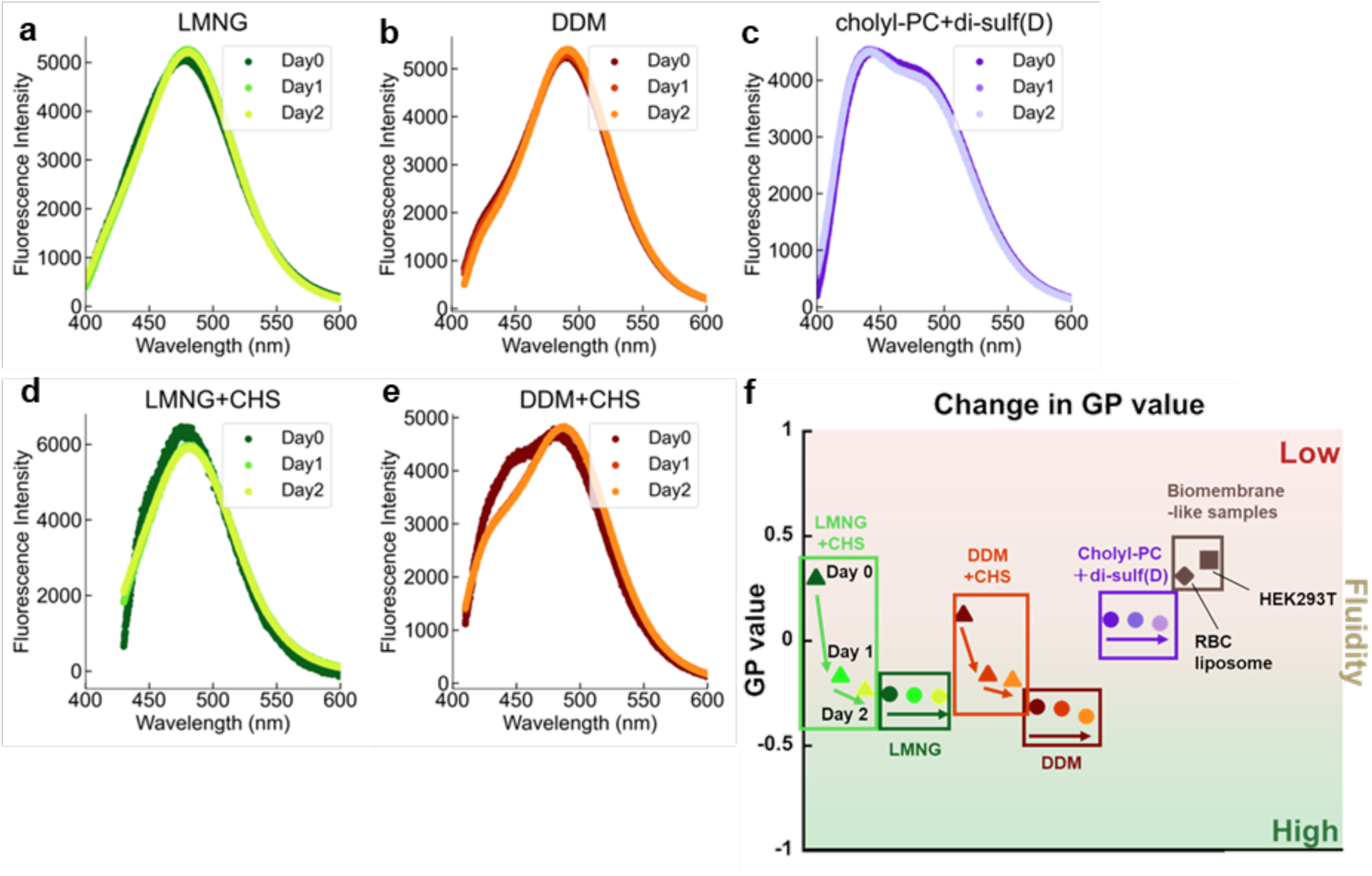
Evaluation of micelle fluidity using Laurdan. **(a-e)** Fluorescence spectra of Laurdan in each detergent micelle on Days 0–2. The spectra are shown after subtracting the background signal of Laurdan-free samples and normalizing it by the integrated area. **(f)** Summary of time-lapse changes in GP values calculated using Eq. (4).

Micelle aggregation was evaluated based on FRET between empty micelles containing donor lipids (lissamine rhodamine B-labeled phosphatidylethanolamine) and acceptor lipids (Atto647N-labeled phosphatidylethanolamine) (Fig. S3a). As a FRET positive control, liposomes containing both donor and acceptor lipids in the same vesicle were used in this experiment. The FRET signal (donor quenching and acceptor-sensitized emission) was observed for the positive control (Fig. S3b), whereas no FRET signals were observed for any micelles stored for 7 days at 25 °C (Fig. S3c, d). The positive control exhibited FRET efficiency, which was defined as the ratio of acceptor (670 nm) to donor emission peaks (590 nm) (Fig. S3b), at a relatively higher level (∼2.5) during observation, whereas those of micelles showed lower levels (∼0.25) than that of the positive control (Fig. S3e and Table S1). Protein aggregation in micelles was analyzed based on changes in the size of the EYFP fluoresce bright spots captured by TIRFM (Fig. S4). The solubilized β_2_AR-EYFP-AviTag was immobilized on the glass substrate to compare the size reflecting protein aggregation each day. We defined bright spots larger than a circle with a radius of 229 nm based on the Rayleigh criterion as aggregations occurring during incubation at 25 °C (Fig. S4a). However, the ratios did not change from Day 0 to Day 2 (Fig. S4b). These data demonstrated that micelle or protein aggregation did not occur, and the change in the ligand-binding ability of β_2_AR-EYFP-AviTag (Fig. 3h) was not due to the surface properties of the micelles.

The lateral pressure within the micelles was analyzed based on the generalized polarization (GP) value, which was calculated from the fluorescence spectrum of the environment-sensitive fluorescent probe, Laurdan^36^. The GP value is an index of the fluidity of lipid membranes; a high GP value indicates low membrane fluidity, whereas a low GP value indicates high membrane fluidity^37,38^. In this experiment, empty micelles were used because it has been reported that the Laurdan spectrum responds exclusively to changes in the lipid environment, even when a membrane-inserting peptide is added at high concentrations^39^, suggesting that the presence or absence of proteins within micelles has a negligible effect on the Laurdan spectrum. Moreover, the content of β_2_AR-EYFP-AviTag varied depending on the type of detergents, as revealed by fluorescence correlation spectroscopy (FCS), (Fig. S5), may complicate the interpretation complicated. Laurdan was solubilized in micelles and the GP values calculated based on Eq. (4) using the fluorescence intensities at 440 and 490 nm^36,40^ in the emission spectra (Fig. 4a–e) are shown in Figure 4f. HEK293T cells and liposomes mimicking the lipid composition of red blood cell (RBC) membranes (hereafter referred to as RBC liposomes) were evaluated^29,41^. HEK293T cell membranes were chosen because it is the native environment for the target β_2_AR, and RBC liposomes were selected because they are widely used as a biological environment model for evaluating membrane physical properties. The lipid composition used for the RBC liposomes in this study was rich in cholesterol and sphingomyelin, which is considered to replicate the liquid-ordered phase present in biological membranes. LMNG and DDM micelles exhibited lower GP values (relatively higher fluidity) than the control biological membranes, whereas the micelles of cholyl-PC + di-sulf(D) showed higher GP values (relatively lower fluidity), which were close to that of the biological membranes, than those of LMNG and DDM (Fig. 4f). No obvious changes in GP values were observed for these detergents from Day 0 to Day 2 (Fig. 4f). These results indicate that the lateral pressure within micelles differs for each detergent and may influence the ligand-binding ability of β_2_AR-EYFP-AviTag for long-term storage (Fig. 3h). In a biological membrane environment, the membrane consists of a mixture of regions with low and high membrane fluidity, and membrane proteins randomly diffuse on the biological membrane to exert their functional activities (Fig. 5). However, solubilized membrane proteins in detergents may have gradually lost their activity by being confined to a specific microenvironment (Fig. 5) (refer to the *Discussion* section for details).

**Fig. 5.**
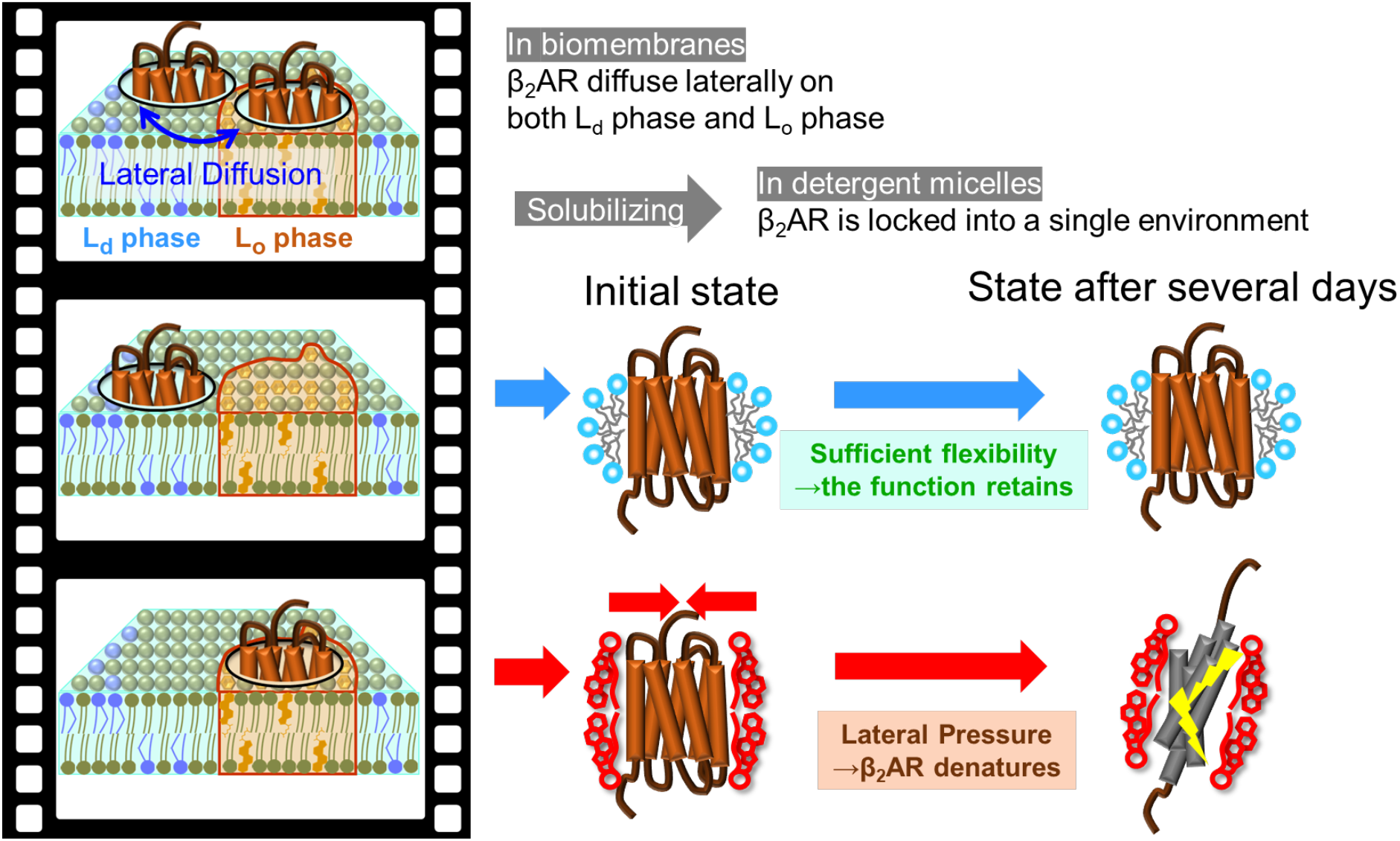
A proposed model to explain the difference in the behavior of β_2_AR in biological membranes and detergent micelles. Membrane proteins randomly diffuse on biological membranes. This figure depicts a frame-by-frame animation of the dynamic diffusion between the L_o_ and L_d_ domains. The microenvironment of the detergent can be described as a snapshot taken from a part of the animation, frame by frame. Therefore, the solubilized membrane proteins in detergents may maintain their native structure and functional activity for a long term if the microenvironment that detergents provide is suitable for them, and *vice versa*, membrane proteins may lose them earlier.

To identify the factors contributing to the differences in GP values between the mixture of cholyl-PC + di-sulf(D) and LMNG or DDM, we focused on the cholesterol backbone of cholyl-PC and di-sulf(D) (Fig. 1d). Cholesterol is widely known to stiffen biomenbranes^42^, which could have affected the results. To prove this hypothesis, cholesteryl hemisuccinate (CHS), which has a steroid backbone similar to the cholesterol structure and a function to stabilize membrane protein and micelle complexes^43–45^, was supplemented with the LMNG and DDM micelles. As expected, the LMNG + CHS and DDM + CHS micelles exhibited higher GP values (relatively lower fluidity) than those of LMNG and CHS alone, which were similar to those of cholyl-PC + di-sulf(D) on Day 0 (Fig. 4f). However, their GP values decreased drastically on Days 1 and 2, returning to their original levels (Fig. 4f). This is probably because CHS-containing micelles temporarily create a microenvironment with low fluidity immediately after preparation, but CHS may rapidly dissociate from the micelles because CHS is not covalently bound to the detergents, and their properties differ from each other.

## Discussion

In this study, we established an evaluation system for the ligand-binding ability of solubilized full-length β_2_AR using TIRFM. By utilizing this system to analyze the colocalization of solubilized β_2_AR and a fluorescent ligand, we successfully and quantitatively observed differences in the stability of β_2_AR depending on the type of detergents. We found that β_2_AR solubilized in cholyl-PC + di-sulf(D) mixed micelles exhibited a drastic decrease in specific ligand binding from Day 1 onward, whereas β_2_AR solubilized in LMNG or DDM micelles relatively retained its ligand-binding ability for several days at room temperature.

This system allows for the quantification of ligand binding using small amounts of unpurified samples and has the potential to become the first-line method for functional evaluation in the following cases: (*i*) membrane proteins with low expression levels^46^, (*ii*) misfolded proteins due to gene mutations and so on causing congenital diseases (*e.g.,* multiple-transmembrane proteins including GPCRs with complicated structures)^47,48^, and (*iii*) membrane proteins with high cell toxicity if overexpressed (*e.g.,* solute transporter transporters)^49^. Focusing on these untouched membrane proteins potentially lead to further expansion of the drug discovery market and address niche demands by overcoming hurdles in research techniques.

To discuss the reason for the difference in the stability of solubilized β_2_AR in detergents, we applied this case to the context of a biological membrane environment. Biological membranes are organized by a wide variety of lipids and proteins, and the heterogeneity generated by complicated interactions between the membrane components causes transient phase separation or domain formation to occur instantaneously^50^. Saturated and unsaturated phospholipid species or cholesterol can be separated into two distinct liquid phases. As shown in Figure 5, a relatively packed, ordered phase enriched in saturated lipids and cholesterol is termed the liquid-ordered (L_o_) phase^51^, and a more fluid, disordered phase comprising mainly unsaturated lipids is termed the liquid-disordered (L_d_) phase^52,53^. Due to its tight molecular packing and enrichment of sterol species and saturated lipids^54^, the L_o_ phase tends to form lipid rafts^50^. Generally, membrane proteins are reported to diffuse laterally between the L_o_ and L_d_ phases on biological membranes to exert their functional activity^55,56^; β_2_ARs translocate to low-fluidity regions only during signal transduction activation^57^. Because detergents provide membrane proteins with a unique microenvironment, similar to that created by isolating a single compartment from the biological membrane (Fig. 5), we considered that β_2_AR with a relatively flexible structure may gradually lose their structural flexibility when confined to a specific microenvironment in micelles, potentially leading to a disruption of its ligand-binding pocket space and a decline in its ligand-binding ability (Fig. 3h). This could be due to the difference in the structural features of the detergents; cholyl-PC + di-sulf(D) contains a steroid backbone that is absent in LMNG and DDM (Fig. 1d), resulting in a difference in fluidity (Fig. 4f)^58,59^. This hypothesis was proven by obtaining higher GP values when CHS was added to LMNG and DDM micelles (Fig. 4f). Although the apparent fluidity of cholyl-PC + disulf(D) micelles was consistent with the overall fluidity of biological membranes, the interior of the cholyl-PC + di-sulf(D) micelle could be a more rigid environment than a biological membrane because the GP values also change depending on the membrane curvature^60–62^. This is presumably because the micelle size of cholyl-PC + di-sulf(D) (∼5 nm)^11^ is much smaller than that of RBC liposomes (several hundred nanometers) and HEK293T cells (approximately 10–15 µm) (data not shown), which presumably allows easier access of water molecules to the micelle interface, resulting in the apparent GP values not reflecting the true fluidity. However, we intend to emphasize that there are optimal detergents depending on the type of membrane proteins because the mixture of cholyl-PC + di-sulf(D) was found to be compatible with membrane proteins, such as bacteriorhodopsin and M2 proton channel of influenza A viruses with relatively rigid structures without any interior free spaces, and successfully maintained their functions and secondary structures for 7 days at 40°C^11^.

Until two or three decades ago, determining the crystal structures of GPCRs was technically challenging; therefore, the development of mutagenesis, chimera fusion, and truncation techniques contributed significantly to advances in crystallization and structural analysis for preventing protein degradation after isolation from biomembranes^2,4,8,9,63–65^. The advent of cryo-electron microscopy has also led to further dramatic advancements in structural analysis techniques^66–68^. In this context, both detergents and lipid nanodiscs have been widely used in many studies. Our analytical method is potentially applicable for screening detergents or lipid nanodiscs suitable for target membrane proteins. However, there is concern that mutated, fused, or truncated proteins may exhibit different three-dimensional structures and functions compared to their native wild-type forms. For example, mutated β_2_ARs have been shown to exhibit slightly different activities when expressed in living cells^32^. It is desirable to analyze the natural structure of full-length membrane proteins without any modifications; however, reports on this issue are still limited^69–71^. Although the C-terminus of β_2_AR was modified in this study to allow for immobilization on the glass substrate, this approach differs from previous methods in that the full-length β_2_AR without any modifications to the core sequence was used for the analysis. Furthermore, this method allows the use of small sample amounts (∼10 ng of β_2_AR-EYFP-AviTag estimated from FCS measurements shown in Fig. S5) without the need for purification, offering advantages in terms of cost and time. Further validation is required to determine whether this method can be applied to GPCRs other than β_2_AR. By comparing the insights accumulated from previous receptor studies with our current results, in the future work, we will investigate the relationship between the functional activity of membrane proteins in their solubilized state and the microenvironment in which they are situated, and ultimately extrapolated these implications to other receptors.

## Conclusion

In this study, we established an analytical method for selecting suitable detergents to maintain GPCR functions. Mammalian β_2_AR was selected as a typical class-A GPCRs because of their relatively flexible structure and ease of denaturation. We evaluated the preservation effects of detergents on receptor functionality for several days at 25 °C by applying TIRFM to track the colocalization of a fluorescent ligand with β_2_ARs immobilized on a glass substrate. Notable features of our analytical methodology are that it allows us to detect specific ligand-binding to solubilized β_2_ARs under unpurified conditions using a small sample volume and distinguish between nonspecific and specific ligand binding. We found that the microenvironment of detergents, such as the lateral pressure within micelles, affects the ligand-binding ability of β_2_AR. LMNG and DDM exhibited a fluid environment and maintained the ligand-binding ability of β_2_ARs for several days after solubilization. In contrast, the mixture of cholyl-PC + di-sulf(D) provided a rigid micellar environment and caused β_2_AR to lose its ability within a few days. Membrane proteins may gradually lose their activity while being confined to an unsuitable state for their functionality. Our methodology allows for the rapid identification of the optimal detergent for target membrane proteins.

## Associated Content Supplementary Information

The Supporting Information is available free of charge at https://XXXX.

Experimental details, receptor constructs, NPY_2_R expression and ligand binding, micelle aggregation, and solubilized protein aggregation, and FCS measurements (PDF).

## Authorship Contributions

**H.H.**: Conceptualization, Investigation, Methodology, Data curation, Formal analysis, Writing – original draft and editing. **K. Kawano**: Conceptualization, Resources, Project administration, Investigation, Methodology, Data curation, Formal analysis, Writing – review and editing, Visualization, Supervision. **T.F., Y.Y., Y.K., K. Kurokawa, K.N., and K.T.**: Resources, Writing – review, and editing. **A.T.**: Methodology, Writing – review and editing. **K.M.**: Conceptualization, Resources, Funding acquisition, Project administration, Writing – review and editing, Supervision.

## Notes

The authors declare no conflicts of interest related to this study.

## Acknowledgments

This work was funded by JSPS KAKENHI (24659016, 17K19485, 19K22493, and 23K18184, for K.M.). The authors thank to Prof. Dr. Haruo Ogawa (Kyoto University), Prof. Dr. Yukihiko Sugimoto (Kumamoto University), and Assoc. Prof. Dr. Masaru Hoshino (Kyoto University) for their kind guidance and valuable advice regarding the functional activity assay and fitting analysis of β_2_ARs. English proofreading for this paper was performed using an AI academic writing tool, “Paperpal” (Editage), which is part of the English proofreading grant program supported by Hiroshima University.

