## Supporting Information for "Analytical methodology for unveiling the physical effect of detergents on the function of full-length membrane proteins using a single-molecule tracking system"

### Table of Contents


**Supplementary Materials and Methods** S3

**Supplementary Figures**

Figure S1 The amino acid sequence of the receptors S4–5

Figure S2 $NPY_2R$ expression and its ligand binding in HEK293T cells S6

Figure S3 Evaluation of the micelle aggregation based on FRET S7

Table S1 Summary in the FRET efficiency S8

Figure S4 Evaluation of the solubilized $\beta_2AR$ aggregation S9

Figure S5 FCS measurements of solubilized $\beta_2AR$ S10

**Supplementary References** S11

#### *Supplemental Materials and Methods*

##### FRET measurements

Lipids were prepared at a molar ratio of 90% POPC and 10% cholesterol, hydrated with Bis-Tris buffer, and subjected to five freeze-thaw cycles. Liposomes were then prepared using an extruder equipped with a 100-nm pore size filter. Micelles were prepared following the same procedure used for the Laurdan samples. Fluorescent lipids serving as a FRET pair—Lissamine Rhodamine B-PE (810150, Avanti Polar Lipids, AL, US; Ex: 560 nm, Em: 583 nm) as the donor and Atto647N-DPPE (Atto-TEC; Ex: 646 nm, Em: 664 nm) as the acceptor—were added to the solution simultaneously with the lipids or solubilizers.

The lipid concentration was determined by phosphorus assay (fiske-subbarow method)[1]. It was confirmed that the concentration of both fluorescent lipids was approximately 2 μM when the liposome lipid concentration was 1 mM, or when the detergent was at the concentration used for protein solubilization.

Furthermore, after measuring the fluorescence spectra of the individual samples, the solutions containing the donor and acceptor dyes were mixed at a 1:1 volume ratio, incubated at 25°C, and subjected to repeated fluorescence measurements. The ratio of the fluorescence intensity (arbitrary units) at 670 nm to 590 nm was used as an indicator of the FRET efficiency.

##### Aggregation evaluation via area analysis of bright spots of immobilized $\beta_2$AR

TIRFM image processing was performed using Fiji version 1.54p[2]. To extract only the first frame from each 1,000-frame video dataset, the Make Substack function was used with the Slice set to 1-1. Prior to image binarization, the brightness and contrast were adjusted to 200–700, and Apply LUT was selected. The Threshold tool was then activated, and the image was binarized using the Moment method[3]. For particle analysis, the binarized image was converted to a mask using "Convert to Mask", and the function "Analyze Particles" was used to acquire information regarding the particles of bright spots. A custom macro was used to automate this series of processes.

##### FCS measurements

The solubilized $\beta_2$AR solution was added to a 35 mm glass-bottom dish (Iwaki), and FCS measurements of EYFP fluorescence were performed using a confocal laser scanning microscope, TCS SP8 X FALCON (Leica Microsystems, Wetzlar, Germany) with 488 nm laser excitation at 25°C. For each sample, a 30-second measurement was repeated 10 times. As described previously, the diffusion coefficient and concentration were calculated by fitting the obtained autocorrelation function curves[4]. The structural parameter was determined to be 7.729, based on the average of 10 measurements using Rhodamine B (diffusion coefficient at 25°C: 430 $\mu m^2/s$), a standard fluorescent dye for FCS[5].

#### $\beta_2$AR-EYFP-AviTag

```
  1 MGQPGNGSAF LLAPNRSHAP DHDVTQQRDE VWVVGMGIVM SLIVLAIVFG
 51 NVLVITAIAK FERLQTVTNY FITSLACADL VMGLAVVPFG AAHILMKMWT
101 FGNFWCEFWT SIDVLCVTAS IETLCVIAVD RYFAITSPFK YQSLLTKNKA
151 RVIILMVWIV SGLTSFLPIQ MHWYRATHQE AINCYANETC CDFFTNQAYA
201 IASSIVSFYV PLVIMVFVYS RVFQEAKRQL QKIDKSEGRF HVQNLSQVEQ
251 DGRTGHGLRR SSKFCLKEHK ALKTLGIIMG TFTLCWLPFF IVNIVHVIQD
301 NLIRKEVYIL LNWIGYVNSG FNPLIYCRSP DFRIAFQELL CLRRSSLKAY
351 GNGYSSNGNT GEQSGYHVEQ EKENKLLCED LPGTEDFVGH QGTVPSDNID
401 SQGRNCSTND SLLWARDPPV ATMVSKGEEL FTGVVPILVE LDGDVNGHKF
451 SVSGEGEGDA TYGKLTLKFI CTTGKLPVPW PTLVTTFGYG LQCFARYPDH
501 MKQHDFFKSA MPEGYVQERT IFFKDDGNYK TRAEVKFEGD TLVNRIELKG
551 IDFKEDGNIL GHKLEYNYNS HNVYIMADKQ KNGIKVNFKI RHNIEDGSVQ
601 LADHYQQNTP IGDGPVLLPD NHYLSYQSAL SKDPNEKRDH MVLLEFVTAA
651 GITLGMDELY KGGGSGLNDI FEAQKIEWHE *
```

$\beta_2$AR

EYFP

AviTag

#### $\beta_2$AR-EYFP-SBP

```
  1 MEIAALEKEI AALEKEIAAL EKGGGASGQP GNGSAFLLAP NRSHAPDHDV
 51 TQQRDEVWVV GMGIVMSLIV LAIVFGNVLV ITAIAKFERL QTVTNYFITS
101 LACADLVMGL AVVPFGAAHI LMKMWTFGNF WCEFWTSIDV LCVTASIETL
151 CVIAVDRYFA ITSPFKYQSL LTKNKARVII LMVWIVSGLT SFLPIQMHWY
201 RATHQEAINC YANETCCDFF TNQAYAIASS IVSFYVPLVI MVFVYSRVFQ
251 EAKRQLQKID KSEGRFHVQN LSQVEQDGRT GHGLRRSSKF CLKEHKALKT
301 LGIIMGTFTL CWLPFFIVNI VHVIQDNLIR KEVYILLNWI GYVNSGFNPL
351 IYCRSPDFRI AFQELLCLRR SSLKAYGNGY SSNGNTGEQS GYHVEQEKEN
401 KLLCEDLPGT EDFVGHQGTV PSDNIDSQGR NCSTNDSLLW ARDPPVATMV
451 SKGEELFTGV VPILVELDGD VNGHKFSVSG EGEGDATYGK LTLKFICTTG
501 KLPVPWPTLV TTFGYGLQCF ARYPDHMKQH DFFKSAMPEG YVQERTIFFK
551 DDGNYKTRAE VKFEGDTLVN RIELKGIDFK EDGNILGHKL EYNYNSHNVY
601 IMADKQKNGI KVNFKIRHNI EDGSVQLADH YQQNTPIGDG PVLLPDNHYL
651 SYQSALSKDP NEKRDHMVLL EFVTAAGITL GMDELYKHHH HHHGSGMDEK
701 TTGWRGGHVV EGLAGELEQL RARLEHHPQG QREP*
```

$\beta_2$AR

EYFP

SBP

#### NPY$_2$R-EYFP-AviTag

```
  1 MGPIGAEADE NQTVEEMKVE QYGPQTTPRG ELVPDPEPEL IDSTKLIEVQ
 51 VVLILAYCSI ILLGVIGNSL VIHVVIKFKS MRTVTNFFIA NLAVADLLVN
101 TLCLPFTLTY TLMGEWKMGP VLCHLVPYAQ GLAVQVSTIT LTVIALDRHR
151 CIVYHLESKI SKRISFLIIG LAWGISALLA SPLAIFREYS LIEIIPDFEI
201 VACTEKWPGE EKSIYGTVYS LSSLLILYVL PLGIISFSYT RIWSKLKNHV
251 SPGAANDHYH QRRQKTTKML VCVVVVFAVS WLPLHAFQLA VDIDSQVLDL
301 KEYKLIFTVF HIIAMCSTFA NPLLYGWMNS NYRKAFLSAF RCEQRLDAIH
351 SEVSVTFKAK KNLEVRKNSG PNDSFTEATN VWARDPPVAT MVSKGEELFT
401 GVVPILVELD GDVNGHKFSV SGEGEGDATY GKLTLKFICT TGKLPVPWPT
451 LVTTFGYGLQ CFARYPDHMK QHDFFKSAMP EGYVQERTIF FKDDGNYKTR
501 AEVKFEGDTL VNRIELKGID FKEDGNILGH KLEYNYNSHN VYIMADKQKN
551 GIKVNFKIRH NIEDGSVQLA DHYQQNTPIG DGPVLLPDNH YLSYQSALSK
601 DPNEKRDHMV LLEFVTAAGI TLGMDELYKG GGSGLNDIFE AQKIEWHE*
```

NPY$_2$R

EYFP

AviTag

#### $NPY_2R$-EYFP-SBP

```
  1 MGPIGAEADE NQTVEEMKVE QYGPQTTPRG ELVPDPEPEL IDSTKLIEVQ
 51 VVLILAYCSI ILLGVIGNSL VIHVVIKFKS MRTVTNFFIA NLAVADLLVN    NPY2R
101 TLCLPFTLTY TLMGEWKMGP VLCHLVPYAQ GLAVQVSTIT LTVIALDRHR
151 CIVYHLESKI SKRISFLIIG LAWGISALLA SPLAIFREYS LIEIIPDFEI
201 VACTEKWPGE EKSIYGTVYS LSSLLILYVL PLGIISFSYT RIWSKLKNHV
251 SPGAANDHYH QRRQKTTKML VCVVVVFAVS WLPLHAFQLA VDIDSQVLDL
301 KEYKLIFTVF HIIAMCSTFA NPLLYGWMNS NYRKAFLSAF RCEQRLDAIH
351 SEVSVTFKAK KNLEVRKNSG PNDSFTEATN VWARDPPVAT MVSKGEELFT
401 GVVPILVELD GDVNGHKFSV SGEGEGDATY GKLTLKFICT TGKLPVPWPT    EYFP
451 LVTTFGYGLQ CFARYPDHMK QHDFFKSAMP EGYVQERTIF FKDDGNYKTR
501 AEVKFEGDTL VNRIELKGID FKEDGNILGH KLEYNYNSHN VYIMADKQKN
551 GIKVNFKIRH NIEDGSVQLA DHYQQNTPIG DGPVLLPDNH YLSYQSALSK
601 DPNEKRDHMV LLEFVTAAGI TLGMDELYKH HHHHHGSGMD EKTTGWRGGH
651 VVEGLAGELE QLRARLEHHP QGQREP*                             SBP
```

**Fig. S1 The amino acid sequence of the receptors.** The sequences start from methionine and end at the stop codon (*). $\beta_2$AR (UniProt: P07550), $NPY_2R$ (UniProt: P49146), and EYFP (UniProt: P42212; GFP_AEQVI variants (S65G, V68L, S72A, T203Y)).

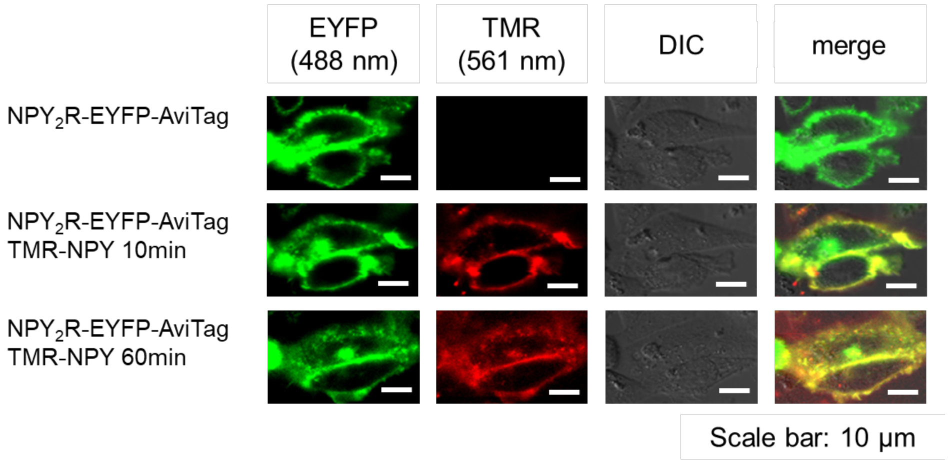


**Fig. S2 $NPY_2R$ expression and its ligand binding in HEK293T cells.**
HEK293T cells expressing $NPY_2R$-EYFP-AviTag in the absence or presence of 100 nM TMR-NPY. The images at 10 and 60 mins after adding TMR-NPY to the cells. At 10 mins, the ligand bound to $NPY_2R$ expressed on the cell membrane was observed. At 60 mins, the internalization of the ligand-bound receptor was observed.

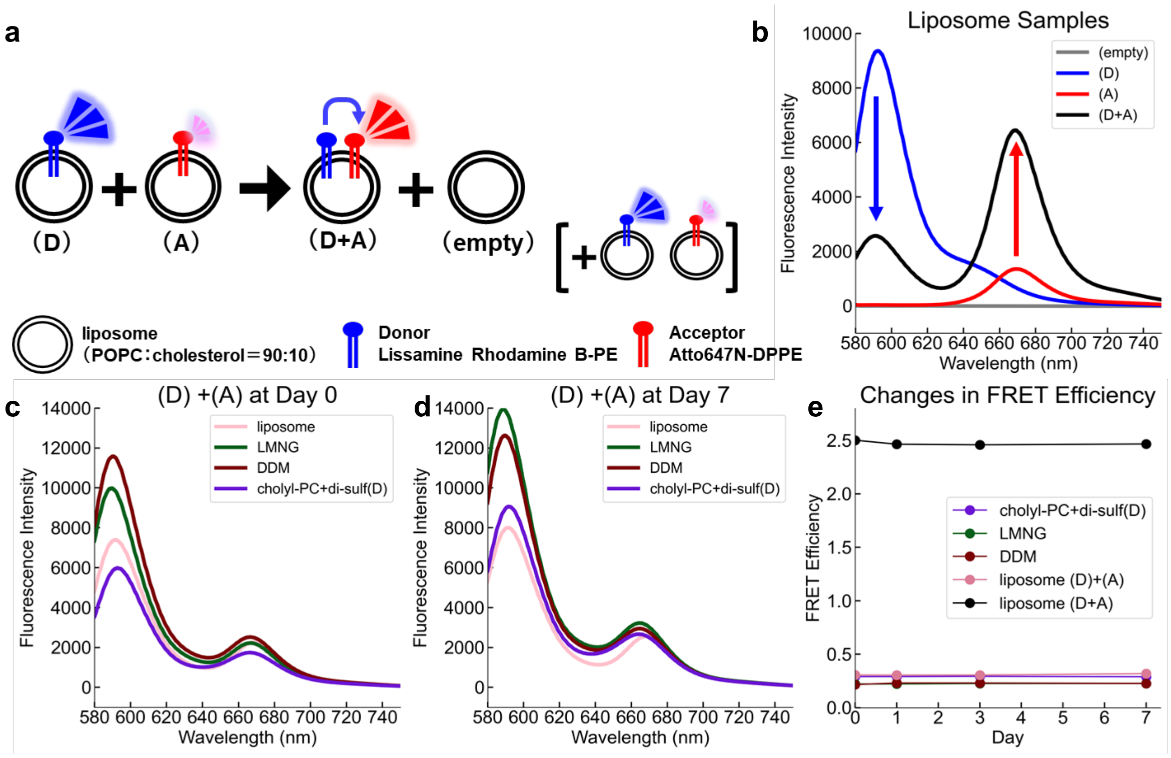


**Fig. S3 Evaluation of the micelle aggregation based on FRET. (a)** A schematic diagram of liposome fusion based on FRET. Lissamine Rhodamine B-PE as the donor and Atto647N-DPPE as the acceptor were used for observing the fusion of liposomes or micelles. **(b-d)** Fluorescence spectra of liposomes or detergent micelles containing donor and acceptor fluorescent lipids. When FRET occurred in the donor and acceptor-ce-existing liposomes (D+A), the donor peak at ~590 nm was diminished, whereas the acceptor peak at ~670 nm increased. When just mixing the donor liposomes (D) and the acceptor liposomes (A), FRET was not observed. **(e)** Time-lapse FRET efficiency. Fluorescence spectra were measured at excitation at 560 nm. The ratio of fluorescence intensities at ~670 nm to that at ~590 nm was used as a FRET indicator.

| Day | LMNG | DDM | cholyl-PC + di-sulf(D) | liposome | liposome (D)+(A) |
|---|---|---|---|---|---|
| Day 0 | 0.291 | 0.221 | 0.216 | 0.306 | 2.50 |
| Day 1 | 0.291 | 0.222 | 0.230 | 0.305 | 2.46 |
| Day3 | 0.293 | 0.226 | 0.230 | 0.305 | 2.46 |
| Day 7 | 0.289 | 0.227 | 0.227 | 0.319 | 2.47 |

**Table S1 Summary in the FRET efficiency.** The FRET efficiency plotted in Fig. S3e are shown here.

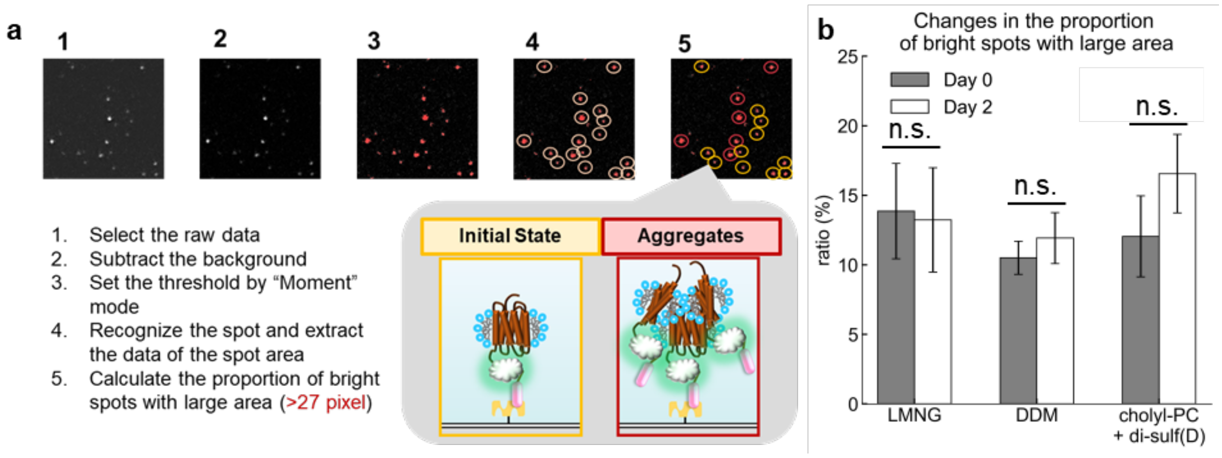


**Fig. S4 Evaluation of the solubilized β2AR aggregation.** **(a)** Outline of the analysis method for spot areas. **(b)** Changes in the proportion of bright spots with large area under each solubilizing condition. Since the TIRFM-based evaluation system established in this study captures moving images of fluorescent spots derived from solubilized β2AR, we reasoned that these image data could be utilized to directly evaluate the aggregation state of the receptor. Image analysis was performed using Fiji, and the Moment method was adopted as the criterion for binarization thresholding[3]. Based on the Rayleigh criterion discussed earlier, the Airy disk diameter of an EYFP fluorescent spot is approximately 450 nm. Given that a single pixel in our TIRFM system corresponds to 78 × 78 nm (6,084 nm$^2$), the theoretical area of the Airy disk spans roughly 27 pixels. Establishing this 27-pixel area as a threshold, we classified spots exceeding this size as multimers/aggregates. We then quantified the ratio of these aggregates to the total spot count, comparing the initial state immediately post-solubilization (Day 0) with the state two days later (Day 2). The analysis revealed a general increase in the aggregate ratio from Day 0 to Day 2 across all β2AR samples, regardless of the detergents. The ratios were not significantly different among all detergents.

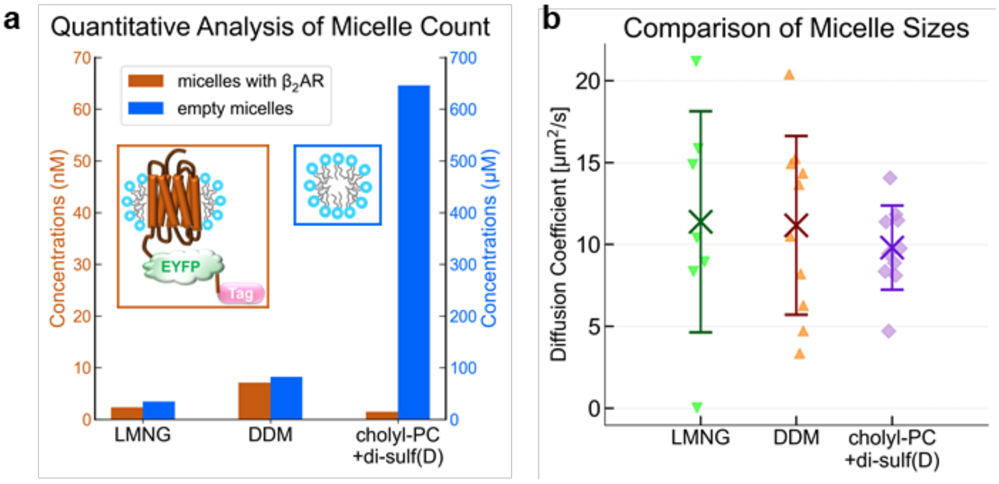


**Fig. S5 FCS measurements of solubilized β$_2$AR. (a)** Concentrations of β$_2$AR-containing micelles (orange, left y-axis) and empty micelles (light blue, right y-axis) for each detergent. The concentration of β$_2$AR-containing micelles was calculated from FCS measurements, while the concentration of empty micelles was estimated based on their lipid concentrations determined by phosphorus assay (fiske-subbarow method). **b.** Diffusion coefficients of β$_2$ARs solubilized in detergents. The cross markers and error bars indicate the mean ± S.D., $n$ = 10.
